# Population level mitogenomics of the long-lived Greater Mouse-eared bat, *Myotis myotis*, reveals dynamic heteroplasmy and challenges the Free Radical Theory of Ageing

**DOI:** 10.1101/224592

**Authors:** David Jebb, Nicole M. Foley, Conor V. Whelan, Frédéric Touzalin, Sebastien J. Puechmaille, Emma C. Teeling

**Affiliations:** School of Biology and Environmental Science, University College Dublin, Belfield, Dublin 4, Ireland; Applied Zoology and Conservation, Zoological Institute, Greifswald University, Greifswald, Germany; Laboratoire Evolution et Diversité Biologique, Université Toulouse 3, Paul Sabatier 31062 Toulouse Cedex 09, France

**Keywords:** Longevity, Mitochondria, Bats, Mutation, Oxidative stress

## Abstract

**Background:** Bats are the only mammals capable of true, powered flight, which drives an extremely high metabolic rate. The “Free Radical Theory of Ageing” (FTRA) posits that a high metabolic rate leads to mitochondrial heteroplasmy and the progressive ageing phenotype. Contrary to this, bats are the longest lived order of mammals despite their small size and high metabolic rate. To investigate if bats exhibit increased mitochondrial heteroplasmy with age as predicted by the FRTA, we performed targeted, deep sequencing of mitogenomes and measured point heteroplasmy in wild, long lived *Myotis myotis* as they age.

**Results:** In total, blood was sampled from 195 individuals, aged between <1 and at 6+ years old, and whole mitochondria were sequenced, with a subset sampled over multiple years. The majority of heteroplasmies, 77.6%, were at a frequency below 5%. Oxidative mutations were not the primary source of heteroplasmies and present in only a small number of individuals, likely representing local oxidative stress events. There was a significant positive correlation between age and heteroplasmy, with a rate of increase of 0.13 sites per year. Longitudinal data from recaptured individuals show heteroplasmy is dynamic, and does not increase uniformly over time.

**Conclusions:** We show that bats do not suffer from the predicted, inevitable increase in heteroplasmy which underscores the FRTA. Most heteroplasmies were at low frequency and are primarily transitions. Heteroplasmy increased with age, however how this contributes to ageing is unclear, as heteroplasmy was dynamic, questioning its presumed role as a primary driver of ageing.

## Background

The mitochondrion in mammals harbours a ~16.5kb, circular chromosome, or mitogenome. Across vertebrates the mitogenome shows a conserved gene content and synteny, containing 13 protein coding genes, 2 ribosomal RNA genes, 22 transfer RNA genes and a control region. Protein coding genes encode essential subunits of the electron transport chain (ETC) complexes, while the tRNA and rRNA genes encode components of the mitochondrial translation machinery. The mitogenome is maternally inherited, is nonrecombining and has a higher mutation rate than the nuclear genome [1, 2]. These properties have made it an attractive study locus for phylogenetics, population genetics and forensics. The mitochondria have also been the focus of ageing research since the 1950s due to their central role in the “Free Radical Theory of Ageing” [3].

The Free Radical Theory of Ageing (FRTA) states, through normal mitochondrial function, reactive oxygen species (ROS) are generated which damage biomolecules leading to mitochondrial dysfunction [3]. ROS are generated when electrons leak from the ETC and reduce oxygen in the absence of hydrogen cations. The mitogenome was once thought to be particularly susceptible to oxidative damage due to its lack of protective histones and close proximity to the ETC, though it is now known that the mtDNA is not naked. Rather, the mtDNA and the TFAM protein compact to form DNA-protein aggregates, called nucleoids, which in mammals typically contain a single mitogenome [2, 4, 5]. These oxidative mutations become hardcoded in the mitogenome, leading to production of mutated ETC components, and mitochondrial dysfunction. Increased mitochondrial dysfunction gives rise to greater ROS production, which cause greater dysfunction in a “vicious cycle”. This gradual accumulation of heteroplasmy and dysfunctional mitochondria is thought to give rise to the familiar progressive, ageing phenotype [5–7].

Mitochondrial heteroplasmy is the presence of multiple non-identical mitogenomes in a single individual [8]. Mitogenomes may vary in length or at single nucleotides, referred to as length and point heteroplasmy respectively. Heteroplasmy plays an important role in mitochondrial disease, cancer and ageing [7, 9–11]. Once thought to be extremely rare in human populations, recent next generation sequencing efforts have found extensive heteroplasmy in healthy individuals [12]. These heteroplasmies are predominantly at low frequency, enriched for pathogenic mutations, and show tissue and allele specific patterns [13–15]. As the mitochondrial genome is maternally inherited, studies of mother-child pairs have shown that maternal age at conception is positively correlated with levels of heteroplasmy [16]. More generally, levels of heteroplasmy have been shown to increase with age in humans, and contribute to debilitating age related diseases, as well as developmental disorders such as autism [10, 13, 15, 17].

The correlated increase in low level heteroplasmies with age in humans seems to support the FRTA. However, human studies have shown a lack of oxidative transversions but instead show transitions are the primary source of mtDNA mutations [13, 15, 18]. Apart from humans, little is known about low level heteroplasmy in other mammals. Studies involving mutant mouse models have been used to test the relationship between of mitochondrial mutation and age. POLG^−/−^ mice express a mitochondrial polymerase with no proof reading activity [19, 20] and so accumulate mitochondrial mutations faster than a wild type mouse. These mice show a strong progeroid phenotype, supporting a role for mitochondrial mutations in ageing [20, 21]. Mammalian studies have been restricted to the control region, often in repeat containing loci, and have not investigated any potential relationship with age [22–29]. Moreover, few comparative studies have taken advantage of recent advances in next generation sequencing technologies. Recently, Rensch et al., (2016) [1] used publically available ChIP-Seq data from 16 species to evaluate patterns of heteroplasmy across the vertebrate phylogeny and concluded that divergent species show similar patterns in frequency and location of heteroplasmy. This study discovered relatively few heteroplasmies, likely due to the high frequency cut off used, and few individuals available for some species. They did not investigate change in heteroplasmy levels with age. To date, next generation sequencing methods have not been used to investigate the relationship between heteroplasmy and age outside of humans.

Chiroptera, the bats, display exceptional longevity for their body size and metabolic rate. Bats are the longest lived order of mammals, living almost ten times longer than expected given their body size [30, 31]. The longest lived bat is *Myotis brandtii* (weight ~7g), which was first captured as an adult and recaptured 41 years later showing negligible signs of senesce [32, 33]. Interestingly, bats are the only mammals capable of true powered flight [34, 35] This energy intensive form of locomotion requires an extremely high metabolic rate. During flight the metabolic rate in bats is up to 3 times higher than that of a terrestrial mammal of the same size during exercise [36]. Bats can increase oxygen consumption 20-30 fold during flight, and *M. velifer* has been shown to increase oxygen consumption rate-130 fold [36, 37]. According to the FRTA, increased oxygen consumption as a result of flight should lead to increased ROS production, thereby accelerating the ageing process. However, bats seem to defy the FRTA [30, 38]. *M. lucifugus* has been shown to produce less ROS per unit of oxygen consumed compared to mammals of a similar size, suggesting *Myotis* bats have increased “mitochondrial efficiency” and have potentially evolved some mechanism to avoid or manage flight-induced oxidative stress [39].

Heteroplasmy has previously been studied in the long lived *Myotis* genus. Petri et al., (1996) [23] studied length and point heteroplasmy in the control region of the *M. myotis* mitogenome. They found high levels of diversity within individuals at a tandem repeat between the tRNA Proline and the conserved sequence blocks of the control region. Sequence diversity at this locus within individuals was similar to diversity between colonies from Germany and Portugal. The length of the repeated array also varied at this locus between and within individuals, with approximately 40% of individuals showing at least two different size arrays. Kerth et al.,[40] studied the homologous repeat array in *M. bechsteinii* from Germany and exploited the high levels of sequence and length diversity in this array to study “microgeographic” population structuring in these bats. This locus has been suggested to possibly regulate mitochondrial transcription due to homology with termination-associated sequences leading to a complex secondary structure [29, 41]. However, this repeat locus is found only in the Vespertilioninae and Myotinae subfamilies [41, 42], and so cannot apply generally to exceptional longevity across the chiropteran phylogeny. Heteroplasmy has also been studied outside the Vespertilionidae. The distantly related *Rhinolophus sinicus* has was found to exhibit point and length heteroplasmy. As in the studies in vespertilionid bats, heteroplasmy was observed at a tandem repeat in the control region using PCR and cloning based techniques [22]. So far, no study has used next generation sequencing to investigate low frequency heteroplasmy across the whole mitogenome in a long lived bat species, or tested if heteroplasmy shows any association with age in bats.

To elucidate if bats exhibit increased mitochondrial heteroplasmy with age as predicted by the FRTA, we performed targeted, deep sequencing of mitogenomes and measured point heteroplasmy in wild, long lived *M. myotis* as they age. We report, for the first time, population level characterization of heteroplasmy based on whole mitogenomes from 195 individuals of known age of the long lived bat, *M. myotis*. Polymerase error likely explains the majority of mutations, with oxidative transversions at a lower rate and concentrated in only a few individuals. Most heteroplasmies are at low frequency, less than 5%. We also sampled and sequenced individuals over multiple years, to assess changes in levels of heteroplasmy at the individual level through time. These unique samples revealed the dynamic nature of heteroplasmy, possibly as a consequence of transient oxidative stress. Contrary to the FRTA, we found that *M. myotis* do not suffer widespread oxidative damage due to flight, instead exhibit acute and transient oxidative stress. We suggest enhanced mitochondrial quality control mechanisms, which may repair or remove damaged mitochondria, possibly contributing to the exceptional longevity in *M. myotis*.

## Results

### Sequencing and Quality Control

252 samples were sequenced obtaining a total of 72,111,278 read pairs prior to quality control. After adapter trimming and stringent quality filtering of reads, 56,608,559 (78.5%) of read pairs were retained. Stringent quality control (QC) resulted in loss of more than 20% of our raw data. 20 samples were deemed unreliable after testing the effect of PCR duplicate removal. Average coverage of samples ranged from 142 to 7800X. 4 samples with average coverage below 1000X were removed. 12 which failed initial QC tests were re-sequenced, and the same quality control steps as before were performed. 4 re-sequenced samples were retained after quality control. A total of 232 samples were retained for further downstream analysis. The average coverage of the 232 QC passed samples was 3837X, ranging from 1355 to 7800X.

### Sample Demography

As the individuals have been tagged with passive integrated transponder (PIT) tags since 2010, it was possible to identify which samples belonged to a unique individual, and know the age for each individual. For individuals caught first as an adult only a minimum bound for the age was known. The 232 QC passed samples included 88 juveniles (<1 years old), and 143 adults (>1 years old) and one individual to which age could not be reliably assigned (Table 1). The 232 samples represent 195 unique individuals, with 21 individuals sampled two, three or four times over consecutive years. All juveniles are 0 years old, with adult samples ranging from 1 years old to at least 7 (hence denoted as 7+). 167 unique individuals aged between 0-5 and a “6+” cohort were used to investigate the relationship between heteroplasmy and age.

**Table 1:**
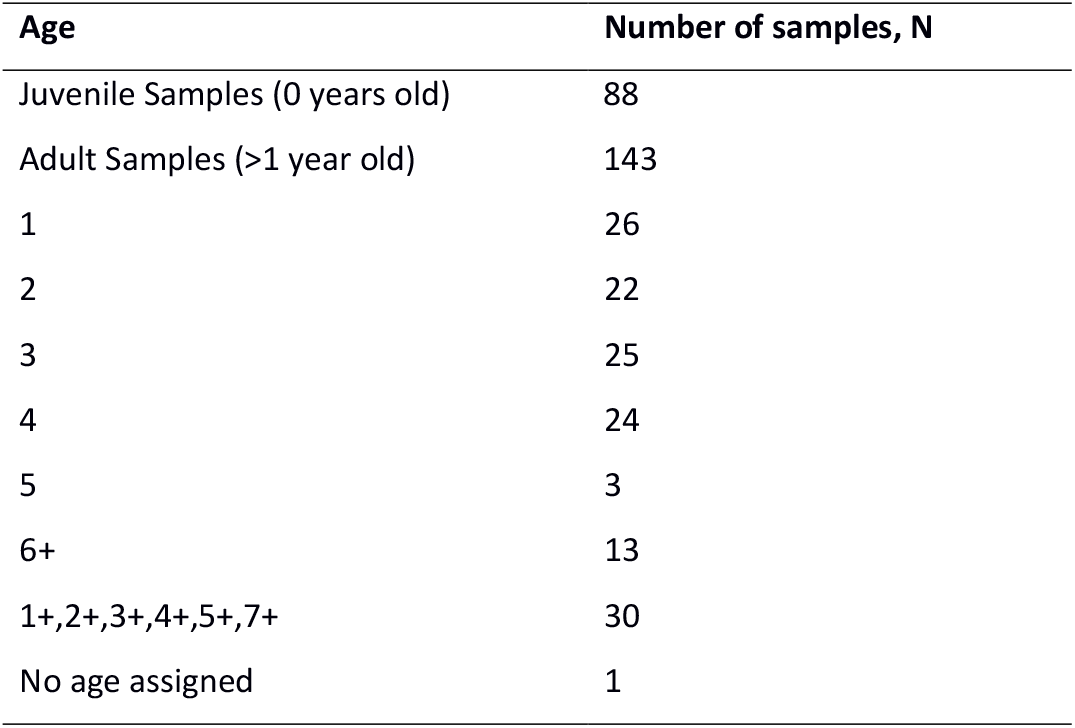
Sample demography

**Table 2:** Analysed Datasets

| **Table 2: Analysed Datasets** |  |  |
| --- | --- | --- |
| **Dataset Title** | **Description** | **Number of samples, N** |
| Sequenced Samples | Total number of samples sequenced | 252 |
| QC Passed | Samples which passed quality control | 232 |
| Unique Individuals | Unique individual bats, recaptures removed | 195 |
| Primary | Unique individuals assigned to age cohorts used in model fitting | 167 |
| Recaptures | 21 individuals caught and sequenced at least twice between 2013 and 2016 | 56 |

## Simulation Results

9 read sets were generated using GemSIM [43], ranging from 50 to 25,000X coverage. Each read set was analysed using one of the three callers LoFreq, VarScan or FreeBayes, producing 27 variant call sets, which were given a score between 0 and 1 defined as (Power*Accuracy)*(1-False Positive Rate) (described in full in Material and Methods, Variant Simulation). LoFreq was the best scoring caller, as shown in Supp. Figure 1, and used to call heteroplasmies in all empirical datasets. FreeBayes consistently had the highest power, detecting more than 85% of true variants even at coverages as low 100X, however, FreeBayes also exhibited a false positive rate >5% at all coverages. LoFreq and VarScan had extremely low error rates, with each detecting only one false positive in the 27 variant call sets. LoFreq had higher power and accuracy than VarScan at all coverage levels (see Additional File 1).

### Frequency and distribution of heteroplasmies in *M. myotis*

From 232 samples, 254 heteroplasmies, across 143 sites in mitogenome, were discovered with a minor allele frequency (MAF) greater than 1%, Figure 1A. The frequency of minor alleles was strongly skewed toward low frequency variants. Figure 1B shows a histogram of number of heteroplasmies by MAF bin for all 254 heteroplasmies. The vast majority, 77.6%, of heteroplasmies are below 5% MAF.

**Figure 1:**
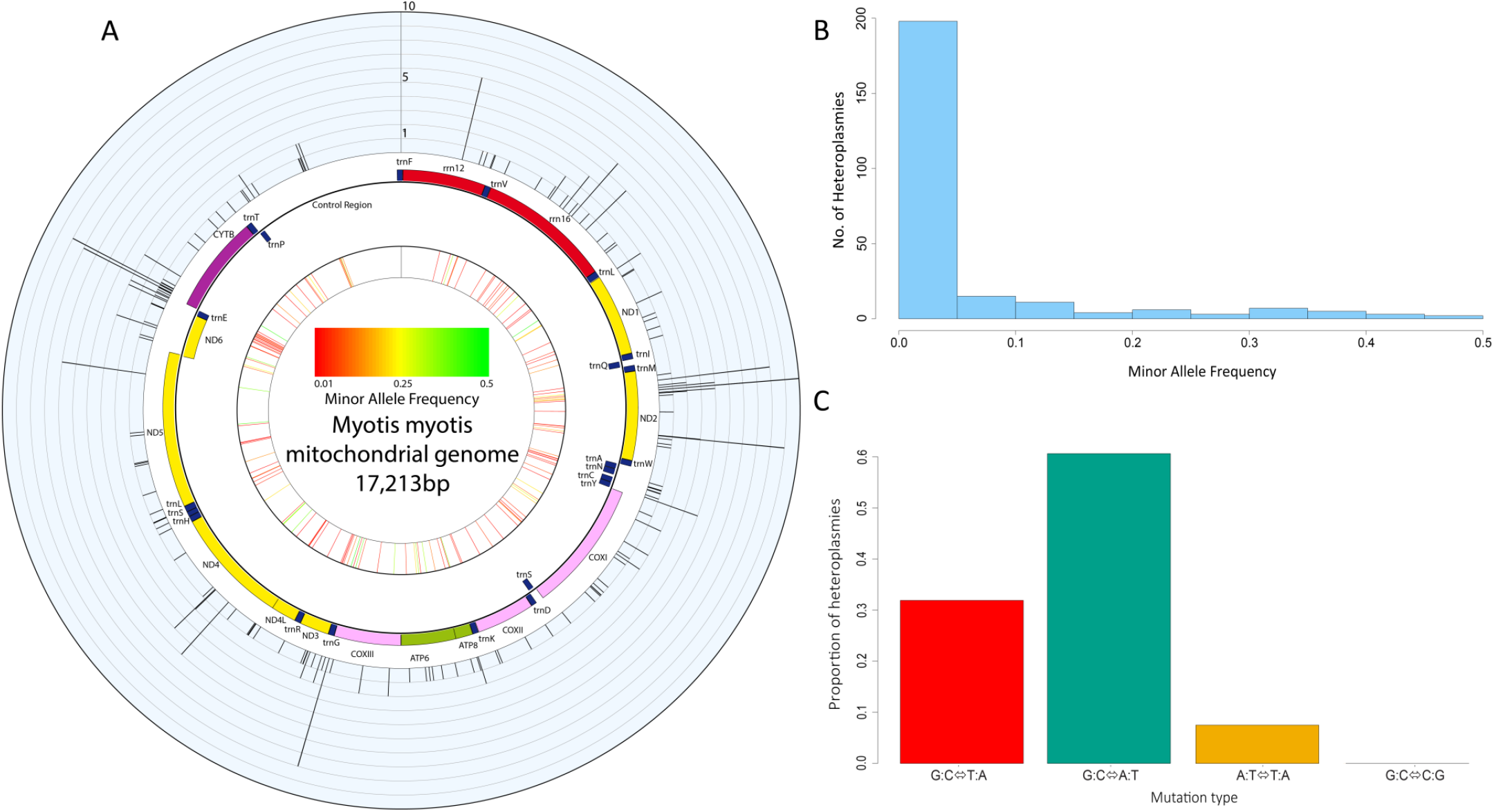
Characteristics of heteroplasmy in *M. myotis*. **A)** A circus plot of the mitogenome of *M. myotis*. In the outer ring histogram, each bar is one of the 143 unique sites which were heteroplasmic in the population, with the height representing the number of individuals sharing the site. The middle ring is a schematic of the mitogenome generated with OGDRAW [81]. The inner most ring is a heat plot showing the average minor allele frequency at each heteroplasmic site, with low frequencies in red and high frequencies in green. **B)** Histogram of minor allele frequencies of 254 heteroplasmies discovered. The vast majority of heteroplasmies were at a frequency below 5%. **C)** Bar chart showing the proportion of mutations attributable to one of four transition or transversion classes. The transition transversion ratio (Ts/Tv) for *M. myotis* was 1.8. Oxidative mutations, G:C⇔T:A transversions, were at a higher level than reported previously in humans.

210 heteroplasmies were collected from unique individuals, using the most recent sample from recaptured individuals. The difference between the frequency distribution of coding and non-coding heteroplasmies was approaching significance (Kolmogorov-Smirnov (KS) Test, p=0.054) with non-coding heteroplasmies tending to be at lower frequencies. Closer investigation of coding heteroplasmies found no significant difference in the frequency distribution of nonsynonymous and synonymous heteroplasmies (KS Test, 0.082), however nonsynonymous heteroplasmies were at lower frequencies than synonymous (Mann-Whitney *U*-test, p=0.021). This also drove the difference observed between coding and noncoding heteroplasmies, as a significant difference was found between the frequency distributions of synonymous and non-coding heteroplasmies, but not between nonsynonymous and non-coding heteroplasmies (KS Test, p=0.006 and p=0.374 respectively) (Supp. Figure 2A). There was also a difference in the frequency of heteroplasmies depending on the codon position with heteroplasmies at the second codon position at a lower frequency than position 1 or 3, which showed no difference to each other (KS Test, p=0.004, p=0.005, p=0.411 respectively) (Supp. Figure 2B). To further explore the effect of heteroplasmies, all nonsynonymous heteroplasmies, excluding nonsense mutations, were scored as “Deleterious” or “Neutral”. Of the 62 nonsynonymous sites scored, 42 were found to be deleterious and 20 neutral. Deleterious heteroplasmies were at significantly lower frequency than neutral heteroplasmies (KS Test, p=0.012).

Heteroplasmic sites were spread relatively evenly across the mitogenome. The number of heteroplasmic *versus* homoplasmic sites were used to construct a contingency table for each mitochondrial partition. Dividing the mitogenome into tRNA, rRNA, protein coding and non-coding regions, showed enrichment for heteroplasmic sites in tRNA genes (p=0.0005, Chi Square Test). Analysis of individual genes revealed *ND2, trnaE* and *trnaN* genes were enriched for heteroplasmic sites (p<0.05, Chi Square Test). The majority of sites, 72.7%, were only observed as heteroplasmic in one individual (Figure 1A), hereafter called “private” heteroplasmies as opposed to “shared”. Private and shared heteroplasmies showed significantly different frequency spectrums (KS Test, p=0.005) with private heteroplasmies at lower average frequency suggesting these were predominantly somatic mutations. Private and shared heteroplasmies showed similar proportions of synonymous and nonsynonymous mutations and no bias toward coding versus non-coding sites (Chi Square test, p>0.1 in all comparisons).

### Transitions are the primary source of heteroplasmies

210 heteroplasmies from 195 unique individuals were classified into 4 classes, 1 transition class (G:C↔A:T) and 3 transversion classes (G:C↔:A, A:T↔T:A and G:C↔:G), as shown in Figure 1C. Transitions were the primary class of heteroplasmy at 60.6%. This was followed by G:C↔T:A transversions at 31.9%, which are characteristic of oxidative stress events. The remaining 7.5% of mutations were A:T↔T:A transversions. *M. myotis* showed a small transition to transversion (Ts/Tv) ratio, with a Ts/Tv of 1.8. Only 38 samples (16.4%) showed evidence of oxidative transversions, of which 33 were unique individuals. The 5 samples containing the highest number of oxidative transversions accounted for 37% of these transversions (30 of 81).

### Correlation between heteroplasmy and age

To test if there was an association with age we fitted a generalised linear model with age as the only explanatory variable, and the count of heteroplasmic sites as the negative binomial distributed response variable. Two samples within the 6+ cohort were found to be unduly influential points using Cook’s Statistic (Supp. Figure 3A and B). As such they were removed and the model fitted again. With these two samples removed no significant association with age was found (ANOVA, p=0.1), as shown in Figure 2. Further, these two samples contained primarily oxidative mutations, one of which contains the highest number of oxidative mutations in our data. To investigate any bias that may be introduced by choosing the most recent sample from recaptured individuals, we produced 1000 permutations in which the sample from recaptured individuals of usable age was chosen at random, to produce a similar dataset of 165 unique individuals. 19.8% of samplings found a significant association between age and number of heteroplasmies (Supp. Figure 4A), with an average slope 0.0969 and ranging from 0.0815 to 0.1237 (Supp. Figure 4B).

**Figure 2:**
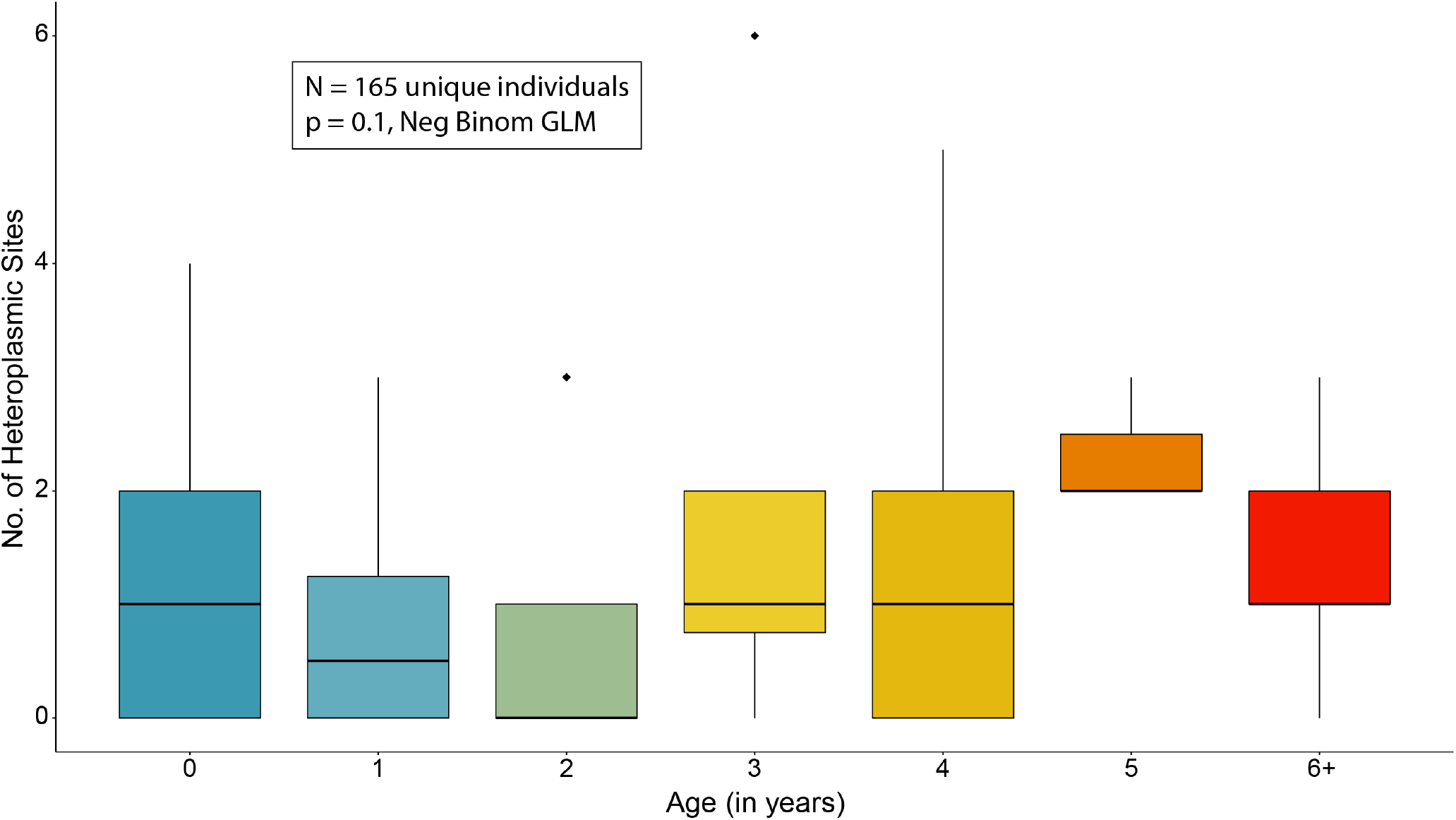
Heteroplasmy is not significantly associated with age in *Myotis myotis*. Boxplots of the distribution of heteroplasmy counts in each age cohort used for age analysis. No significant association was found between age and heteroplasmy using a negative binomial generalized linear model, after the removal unduly influential points.

### Longitudinal analysis of heteroplasmy

From the 27 individuals sampled more than once, we were able to obtain samples from 21 individuals from at least two time-points, which passed quality control. 3 individuals were sampled at 4 time-points, 8 individuals sampled at 3 time-points and 10 individuals at 2 time-points, individuals sampled at 3 or 4 time-points are shown in Figure 3. The mean change in number of heteroplasmies between two consecutive years across individuals was 0.303. Of the 33 consecutive intervals (e.g. 2013 to 2014 or 2014 to 2015 etc.) 15 showed no change, 10 showed a decrease and 8 showed an increase (Supp. Figure 5). Not all heteroplasmies were shared between time-points. 14.7% of heteroplasmies were observed in at least two time-points, with 13% shared between all time-points for an individual. Shared sites ranged in frequency from 43.36% to as low as 1.02%. Some individuals were extremely consistent over all time points. For example individual 000715B9C8, shown in Figure 3, initially showed 2 heteroplasmies, but subsequently lost one. The remaining heteroplasmy was at a low frequency but was found at all 4 time-points and had an extremely consistent frequency, range_maf_ 1.02 – 1.83%. However some individuals were more dynamic. These “spikes” in the number of heteroplasmy was normally followed by a marked decrease the following year. Two bats, 000702E6B0 and 000702FF51 (also shown in Figure 3), showed spikes and subsequent loss, though they retained a subset of the mutations acquired during the spike. Four individuals showing spikes in the number of heteroplasmy, with an increase of 4 or more sites between two subsequent years, were previously found to have some of the highest levels of oxidative mutations. Interestingly, bat 000702EA84 showed a spike in heteroplasmy with little recovery the following year. Instead, this bat may have experienced oxidative stress events during two consecutive field seasons.

**Figure 3:**
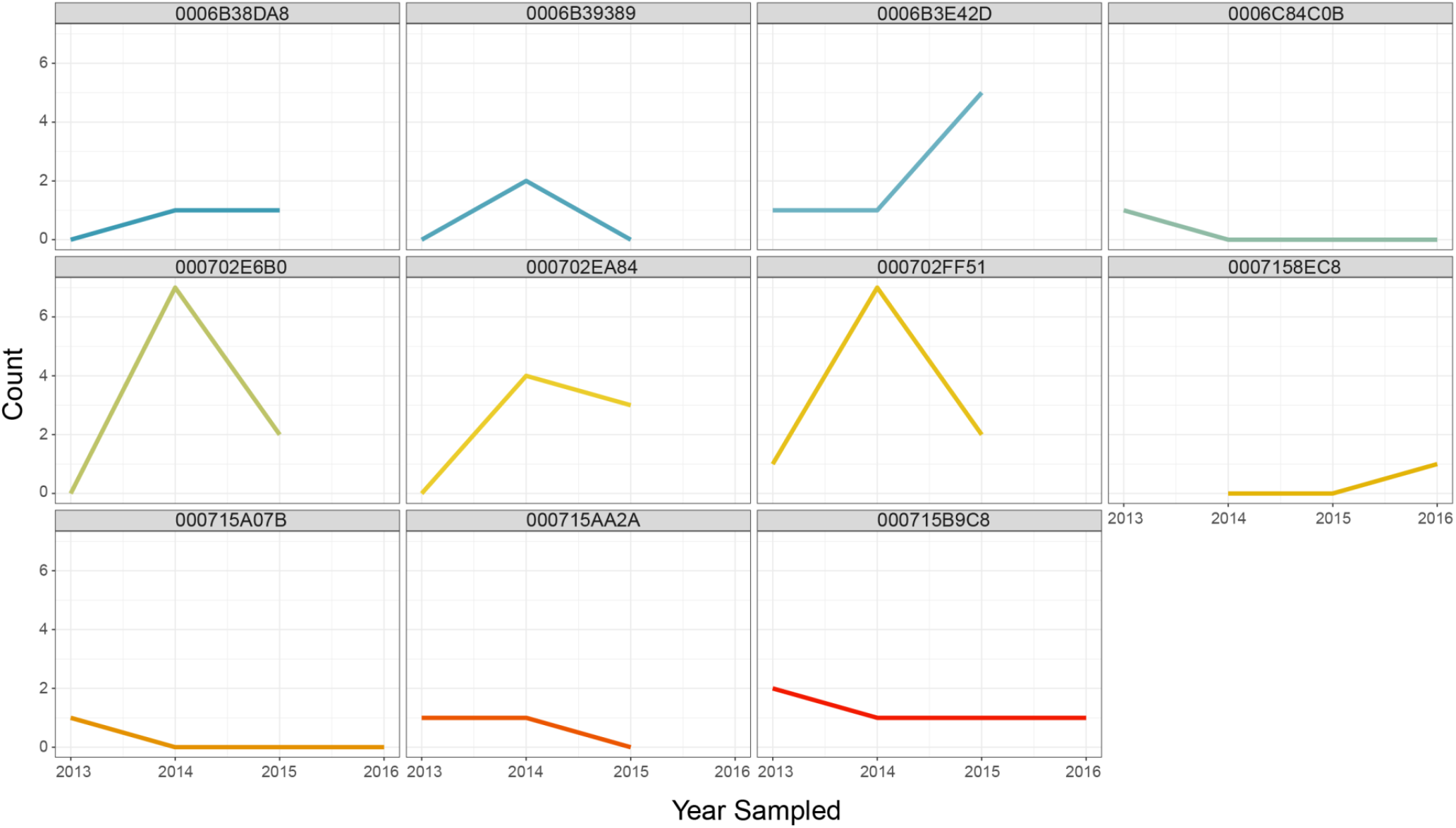
Plots of heteroplasmy count against year sampled for 11 individuals. Heteroplasmy counts through time for 11 individual bats which were sampled 3 or 4 times. Numbers above each plot indicate PIT number used to identify each individual. Some individuals were extremely consistent while others increased sharply, always followed by a decrease where subsequent samples were available.

## Discussion

Comparative genomics has emerged as a powerful new tool in ageing research [30, 44]. However, though mitochondrial dysfunction and heteroplasmy have been long implicated in ageing, there remains a paucity of data from non-human or non-model organisms to associate increased heteroplasmy as a mechanism underpinning the progressive, ageing phenotype. To this end, for the first time we have deep sequenced, to an average depth of >3500X, whole mitogenomes from 195 *M. myotis* individuals, and investigated the association between heteroplasmy and age in these long-lived bats. After outlier removal, we found no evidence for an increase in heteroplasmy with age. We also found little to no evidence to support the Free Radical Theory of Ageing, finding no chronic increase in oxidative mutations with age. Through unique, longitudinal sampling we found bats may experience local oxidative stress events followed by removal of the majority of mutations. Nonsynonymous, protein coding mutations were also at a significantly lower frequency than synonymous, and those with predicted deleterious effects were lower again. Together our results suggest *M. myotis* can remove deleterious mitochondrial DNA mutations, preventing their expansion, to maintain mitochondrial homeostasis, and possibly promoting longevity.

The FRTA posits oxidative mutations in the mitogenome should drive mitochondrial dysfunction [3]. However, many studies have shown a dearth of oxidative mutations in mitochondrial DNA, which has led some to question this central tenet of the FRTA [45]. In *M. myotis*, transitions, not oxidative transversions, are the primary source of mutation. Transitions are likely due to errors during DNA replication or spontaneous deamination of nucleosides, rather than damage accrued by oxidative stress [4, 7]. Still, the proportion of heteroplasmies owing to oxidative transversions in *M. myotis* mitochondrial DNA is much higher than reported in humans [13, 15]. However, the majority of these oxidative mutations were concentrated in only a few individuals. These likely represent local oxidative stress events, rather than a population wide phenomenon. The longitudinal data supports this interpretation as two individuals exhibiting high numbers of oxidative transversions, and some of the highest levels of heteroplasmy, were also sampled the following year (Individual 000702E6B0 and 000702FF51, Figure 3). They had cleared or repaired the majority of the oxidative damage, though each retained at least one of the heteroplasmies from the previous year. We suggest *M. myotis* experience local, transient oxidative stress followed by return to homeostasis and removal or repair of oxidative lesions.

As oxidative damage is concentrated, and transient, within a small proportion of individuals, it is unlikely to be the ultimate driver of ageing, as posited in the FRTA. It also seems unlikely that flight is the cause of this oxidative stress, as it should then be ubiquitous across the population rather than concentrated in few individuals. There are multiple other explanations for the transient oxidative damage observed in only some individuals, including infection and recent reproductive effort [46–48].

In recent years, the role of ROS as essential signalling molecules in immune cells has become clear. *In vitro*, macrophages recruit mitochondria to the phagosome where promotion of H_2_O_2_production contributes to the control of *Salmonella* infection [48]. Activation of the NLRP3 inflammasome has also been shown to promote ROS production [47]. It has been suggested that bats have a constitutively active innate immune system, possibly explaining their unique ability to act as reservoirs for diverse viruses, such as SARS, which can be pathogenic in humans and other mammals [49–52]. The constitutive expression of interferon or the presence of circulating viruses in bats may lead to acute levels of ROS production in bats enabling them to quickly mediate infection [53, 54]. The long lived bat species, *Corollia perspicillata*, has also been shown to experience increased oxidative stress during simulated bacterial infection [55]. Potentially, individuals showing increased oxidative damage were experiencing immune related oxidative stress at the time of sampling, although this remains to be tested. This may also suggest that spikes in oxidative transversions are also a blood specific phenomenon; future studies incorporating different tissues may shed light on the role and effect of ROS in the bat immune system.

A second possibility is reproductive effort. It has been shown that reproductive effort increases oxidative stress in free living animals, though not unequivocally [46]. As the *M. myotis* sampled over the course of the study were caught at maternity roosts, most females will have recently given birth to a pup, and may still be lactating. Under the disposable soma theory of ageing, reallocation of resources from self-maintenance to reproductive effort leads to increased damage of somatic tissue during reproductive phases, with unrepaired damage accumulating with time [46, 56]. The spikes in oxidative damage followed by recovery as observed in our longitudinal sampling, showed while most oxidative damage observed during the spike were eventually lost or removed, some mutations persisted, and could be detected the following year. Further longitudinal sampling, or sampling at different life history phases may help elucidate the cause of oxidative stress in these bats.

The relationship between age and heteroplasmy was unclear in bats. A positive correlation with age seen in our data was driven by two samples. These samples were clear outliers with a large effect on our regression. Removal of these samples also removed the association with age. The outliers were both in the oldest cohort and had two of the highest levels of heteroplasmy, primarily due to oxidative mutations. It seems likely these individuals may have been experiencing acute oxidative stress, and may have removed the majority of these oxidative mutations in following years, as seen in other individuals. However, this remains speculative as we have yet to re-sample these two individuals. Further, the observed dynamic nature of heteroplasmy questions the direct contribution of heteroplasmy to ageing in this species. However, as our study population has only been tagged since 2010 we have only sampled a small portion of the life span of these bats, which can live up to 37.1 years. It is possible accumulation of heteroplasmy is not detectable over this time scale. However, if bats aged at a rate predicted by their body size we would expect to see accumulation of heteroplasmy even over this time scale.

The question remains, how do bats maintain their efficient mitochondria, and how might this lead to an increased lifespan? Our data shows no support for flight induced oxidative damage accumulating in *M. myotis*. However, oxidative stress is observed in numerous individuals, and those with samples from subsequent time-points show removal of the majority of mutations, suggesting mitochondrial stress followed by repair or removal of damaged mitochondria. Further, nonsynonymous mutations were at significantly lower frequency than synonymous, suggesting purifying selection acting on these heteroplasmies. Nonsynonymous sites predicted to be deleterious were even lower in frequency again than those with little or no functional impact. This suggests *M. myotis* can prevent expansion of nonsynonymous mutations, which may contribute to the maintenance of mitochondrial function with age.

There are multiple mechanisms by which *M. myotis* may remove and prevent expansion of mitochondrial mutations. One such mechanism is through direct repair of DNA damage. DNA repair genes and pathways have been shown to be evolving or expressed differently in bats compared to other mammals [57–59], however, many of these pathways are not active in the mitochondria. Base excision repair (BER) is the primary method of DNA repair in mitochondria. While double strand break repair operates in mitochondria, the mechanisms have not been fully characterised [4]. Another mechanism by which *M. myotis* may remove oxidative damage is through mitophagy. Mitophagy is a form of autophagy through which cells remove dysfunctional mitochondria. Inhibition of autophagy, and so mitophagy, by knockdown of autophagosome genes has also been shown to lead to increased ROS production [45]. Mitochondria derived ROS have also been shown to promote autophagy during starvation, through oxidation and inactivation of the ATG4 protein [60]. Long lived bats have also been shown to have increased macroautophagy activity in comparison to a short lived, phylogenetically close sister taxa. Interestingly, other homeostatic mechanisms such as chaperone protein levels and proteasome activity, showed no difference in long and short lived bats [61]. This suggests long-lived bats rely on autophagy for maintaining homeostasis. Indeed, previous studies have shown that autophagy genes are upregulated in the blood of *M. myotis* compared to other mammals [59].

Potentially flight may induce selection for efficient mitochondria in bats [62]. *M. lucifugus* juveniles produced more H_2_O_2_ than adults across three tissues; however, this difference was most striking in pre-fledged juveniles. Cloning of the Hyper Variable Region I of the *M. lucifugus* D-loop found less variation in adults compared to juveniles, suggesting convergence on a single mitotype within individuals with age [62]. This may be due to intra-individual selection for efficient mitotypes due to the high energy demands of flight [62]. Endurance training has also been shown to induce mitochondrial health in mutator mice completely rescuing the progeroid phenotype. Mice experiencing an endurance exercise regime showed wild type lifespan, increased aerobic capacity, and indirect evidence for decreased mtDNA mutations [19]. Coupled with our results, this suggests bats may possess stringent mitochondrial quality control to prevent accumulation and expansion of deleterious mutations, maintain mitochondrial function leading to their extraordinary longevity.

## Conclusions

Overall, we found little support for the long standing “Free Radical Theory of Ageing” in the exceptionally long-lived bat, *M. myotis*. Instead, oxidative mutations were not the primary source of mutations and most seemed to be transient, arising from local oxidative stress events before disappearing. We did not find that accumulation of heteroplasmic sites in the mitogenome increases with age in these bats, however, we have only analysed a small portion of the lifespan of this species. Unique longitudinal sampling revealed the dynamic nature of heteroplasmy, with some bats gaining or losing up to 6 heteroplasmies between consecutive years. These drastic shifts in heteroplasmy suggest increases in heteroplasmy may not only be tolerable but removable in *M. myotis*. We propose that stringent mitochondrial quality control mechanisms and intra-individual selection may drive mitochondrial health potentially contributing to the exceptional longevity of *M. myotis*.

## Methods

### Capture and Sampling

All procedures were carried out in accordance with the ethical guidelines and permits delivered by University College Dublin and the Préfet du Morbihan respectively. *M. myotis* were sampled in western France in 2013, 2014, 2015 and 2016 (detailed in Table 3) as described in Huang et al (2016).

**Table 3:** Colony location and sample numbers

| Table 3: Colony location and sample numbers |  |  |  |
| --- | --- | --- | --- |
| Colony | Latitude | Longitude | No. Samples |
| Béganne | N47°35' | W2°14' | 112 |
| Noyal-Muzillac | N47°35' | W2°27' | 20 |
| Férel | N47°28' | W2°20' | 78 |
| La Roche-Bernard | N47°31' | W2°17' | 32 |
| Limerzel | N47°38' | W2°21' | 10 |

## DNA Extraction

Prior to DNA extraction all blood samples had RNA extracted using the RNAzol BD for blood kit (catalogue number RB 192, Molecular Research Centre, Inc.) using the manufacturers protocol with minor modification as per [63]. Post extraction of the RNA containing layer, the phenol phase/interphase was placed at −80°C. DNA was extracted from the phenol phase/interphase using the protocol detailed in the RNAzol BD kit manual with minor modifications. Briefly, for each original volume of blood, the DNA extraction buffer was prepared with 1 volume of 4 M guanidine thiocynate solution, 0.1 volumes of 3 M sodium hydroxide and 0.005 volumes of polyacryl carrier. Samples were allowed to thaw in the DNA extraction buffer and then were vortexed vigorously for 30 s. Samples were incubated at room temperature (RT) for 10 minutes, then vortexed for 30s. 0.2 volumes of chloroform was added to each sample, briefly vortexed, and centrifuged at 12000rpm for 20 mins at RT. The upper fraction was extracted, and extracted once more with an equal volume of chloroform where necessary, to produce a clear, aqueous DNA phase. DNA was precipitated overnight at −20°C in an equal volume of isopropanol. DNA was pelleted by centrifugation at 12000rpm for 20 mins at 4°C. DNA pellets were washed twice with 75% ethanol and then resuspended in 30μl nuclease free water.

### mtDNA Enrichment and Sequencing

The whole mitogenome of *M. myotis* was amplified and sequenced using the primers, and sequencing protocols previously outlined in Jebb et al., (2017)[64]. Briefly, samples were enriched for mitochondrial sequences using long range PCR, in two overlapping ~10kb fragments. Both amplicons were successfully amplified for 252 samples. The two amplicons for each of the 252 samples were purified by vacuum filtration (MilliporeHTS Filtration Plate, cat. no. MAVM0960R), quantified using a nanodrop spectrophotometer and then pooled in equimolar amounts. Pooled amplicons were used to produce sequencing libraries using the Nextera XT preparation kit as per the manufacturer’s instructions. Varying numbers of samples were multiplexed and loaded onto an Illumina MiSeq.

### Quality Control

Raw reads were initially processed using a stringent quality control pipeline prior to variant calling. Samples were trimmed of Illumina and Nextera adapter sequences using Cutadapt (v1.8.3) [65] and The NGS QC Toolkit [66] was used to filter out reads for which 80% of bases had quality scores less than Q30. Due to extreme sequence depths at small target regions in resequencing studies, it is possible that inserts will begin and end at the same position due to chance, called sampling coincidence. Reads from such inserts will be falsely identified as PCR duplicates. However, not removing true PCR duplicates may affect the accuracy of variant calling. In order to remove any potential bias introduced by duplicate removal or retention, we ran the heteroplasmy detection pipeline with duplicates present and removed. As some samples showed extremely high levels of variation we used the interquartile range to identify and remove outliers. An outlier was defined as any value for which the count value was 2 interquartile ranges below the first quartile or above the third quartile. The mean and standard deviation of these filtered counts was then calculated. A cut-off value of two standard deviations from the mean was rounded down to the nearest integer value. Samples with a difference in counts between the two treatments, which was greater than this cut-off were deemed unreliable and removed from downstream analysis. Finally, average coverage was calculated for each sample using the depth command in samtools (v0.1.19) [67]. As all variant callers exhibited power below 80% at coverages below 1000X (Fig. 2), samples with an average coverage below this threshold were removed.

### Heteroplasmy Detection Pipeline

The first and last 500 bp of the mitogenome were copied to the opposite ends, to extend the reference and account for circularity of the mitogenome. Reads were mapped against the extended *M. myotis* mitogenome using the BWA-MEM [68] algorithm. Prior to variant calling BAM files were processed using Picard Tools (v1.90) [69] and the Genome Analysis Tool Kit (GATK) [70]. Briefly, SAM files were sorted and read groups added using Picard Tools. SAM files were then converted to BAM files and indexed. Prior to base quality recalibration a set of raw variants was produced using the chosen low frequency variant caller, either LoFreq Star [71] or VarScan 2 [72]. These raw variants were used as the “known sites” necessary for the BQSR walker in the GATK. As no databases of variants exist for most non-model organisms, these raw variants provide an *ad hoc* solution. Final heteroplasmies were called on the recalibrated data again using the caller of choice. A third variant caller, FreeBayes [73], does not require recalibrated or realigned data, and so these steps were omitted when FreeBayes was used. For the empirical data, variants with a minor allele frequency greater than 1% and at sites with greater than 1000X coverage were retained. These variants are then annotated, removing variants in primer binding sites and the two Myotinae repeats in the control region. An illustration of the bioinformatic workflow for a single sample from quality processing through to annotated variants is depicted in Supp. Figure 6. GNU Parallel (v20170122) [74] was used to parallelise the mapping and processing of SAM and BAM files, markedly decreasing computational time. The number of samples to be run in parallel can be defined by the user.

### Variant Simulation

In order to gauge the sensitivity, accuracy and false positive rate of our bioinformatic pipeline, known variants and Illumina sequence data were generated *in-silico* using the GemSIM package (v 1.5) [43]. Sequence reads in FASTQ format from a reference individual (MMY104, used to generate the reference mitogenome for this species) were mapped to the previously published *M. myotis* mitogenome [64] using the BWA-MEM algorithm, producing a SAM file which was used to produce a general error model using the GemErr.py module. The reference mitogenome was used to generate haplotype information for variant simulation with the GemHap.py module. In total 500 known variants were simulated, in sets of 100 at frequencies of 1%, 4%, 7%, 10% and 15%. The reference mitogenome, error profile and haplotype information were used to generate 250bp paired end Illumina reads. Datasets were simulated at 50X, 100X, 250X, 500X, 1000X, 2500X, 5000X, 10,000X and 25,000X coverage. Predicted variant positions were checked against the known set. Power (proportion of true sites called), false positive rate (proportion of erroneous sites called) and accuracy (proportion of true sites with predicted frequency within ±0.01 of the true frequency) were calculated on each dataset for each caller, FreeBayes, LoFreq and VarScan2 (see Heteroplasmy Detection Pipeline). Finally a score between 0 and 1 was assigned to each caller defined as (Power*Accuracy)*(1-False Positive Rate). All variants called from real samples used the best performing caller.

### Characterising heteroplasmy in *M. myotis*

All filtered and annotated heteroplasmies were collected and manually inspected. Heteroplasmies were binned into several overlapping classes: protein-coding, synonymous, nonsynonymous, non-protein-coding, tRNA, rRNA, non-coding, and by each mitogenome feature. Nonsynonymous mutations were further binned for possible effect as “Deleterious” or “Neutral” based on the PROVEAN algorithm as implemented on the dedicated web server. Default cut-offs were used such that a mutation was binned as “Deleterious” if the computed PROVEAN Score was greater than 2.5 or less than −2.5 [75, 76]. Frequency distributions for each bin were compared using a two tailed non-parametric Kolmogorov-Smirnov Test. The number of unique heteroplasmic sites in a bin was tested for enrichment against the remainder of mitogenome sites analysed (length of mitogenome – primer regions – number of sites cover by bin), using Pearson’s Chi-Square test.

### Correlation between heteroplasmy and age

We constructed a primary dataset of n=167 unique individuals of known age and an oldest cohort, divided into cohorts from 0 to 6+, described in Table 1 and 2, taking the most recent sample of an individual had the bat been sampled multiple times. This dataset was used to test for any significant correlation between number of heteroplasmic sites and age by fitting a generalised liner model (GLM) in R (v3.3.3) using the “MASS” package [77] with age as the sole explanatory variable and number of heteroplasmic sites as the response variable modelled under a negative binomial distribution. A negative binomial distribution was used due the presence of overdispersion in the data making a Poisson distribution unsuitable. The “boot” package [78, 79] in R (v3.3.3) was used to diagnose samples which may be unduly influential points using the Cook Statistic [80]. Influential points were removed and the model was fitted again. To investigate any bias that may be introduced by manually selecting a sample from recaptured individuals, we performed 1000 samplings in which the sample from recaptured individuals of usable age was chosen at random, to produce 1000 datasets (of n=165).

### Longitudinal Analysis

*M. myotis* show natal philopatry, returning to the same roost or broad area to give birth each year. As such, it was possible to recapture the same individuals almost every year. 27 individuals with samples from at least two time-points were sequenced; samples that passed quality control were retained, for a final set of 56 samples from 21 individuals (Tables 2 and 4). We calculated the mean change between any two consecutive years over all individuals, excluding 2 which were sampled non-consecutively. We also calculated the proportion of shared heteroplasmies between any two time-points, and those shared between all time-points.

**Table 4:** Recaptured individuals and associated samples

| Number of times sampled | Number of Individuals | Number of Samples |
| --- | --- | --- |
| 4 | 3 | 12 |
| 3 | 8 | 24 |
| 2 | 10 | 20 |

## Declarations

### Ethics approval and consent to participate

All procedures were carried out in accordance with ethics approval issued by University College Dublin to Prof. Emma Teeling (AERC_1338_Teeling) and in accordance with Arrêté préfectoral permits issued by the Préfet de Morbihan to Frédéric Touzalin and Sébastien J. Puechmaille.

### Consent for publication

Not applicable

### Availability of data and material

Annotated variants and sample information used for statistical analyses are available within Additional File 1. The scripts and binaries for all dependencies needed to run the 3ML pipeline are available at https://github.com/jebbd/3ML. Sequence data generated during the course of the project from real and simulated data are available from the authors on request.

### Authors’ contributions

E.C.T and D.J devised the study. All authors participated in the capture and sampling of *M. myotis* bats. D.J and C.V.W extracted DNA and prepared samples for sequencing. All sequence data was analysed by D.J. D.J was responsible for statistical analyses. D.J is responsible for the figures presented throughout. The manuscript was written by E.C.T. and D.J. with input from all authors.

## Acknowledgements

The authors would like to thank the European Research Council for funding. For assistance in the field we would like to thank: members of Bretagne Vivante, Dr. Serena Dool, students from UCD and all owners/local authorities for allowing access to the sites. We would like to thank the members of the Teeling lab and Dr. John A. Finarelli for helpful discussion of the analyses.

---

**Supplementary Figure 1:**
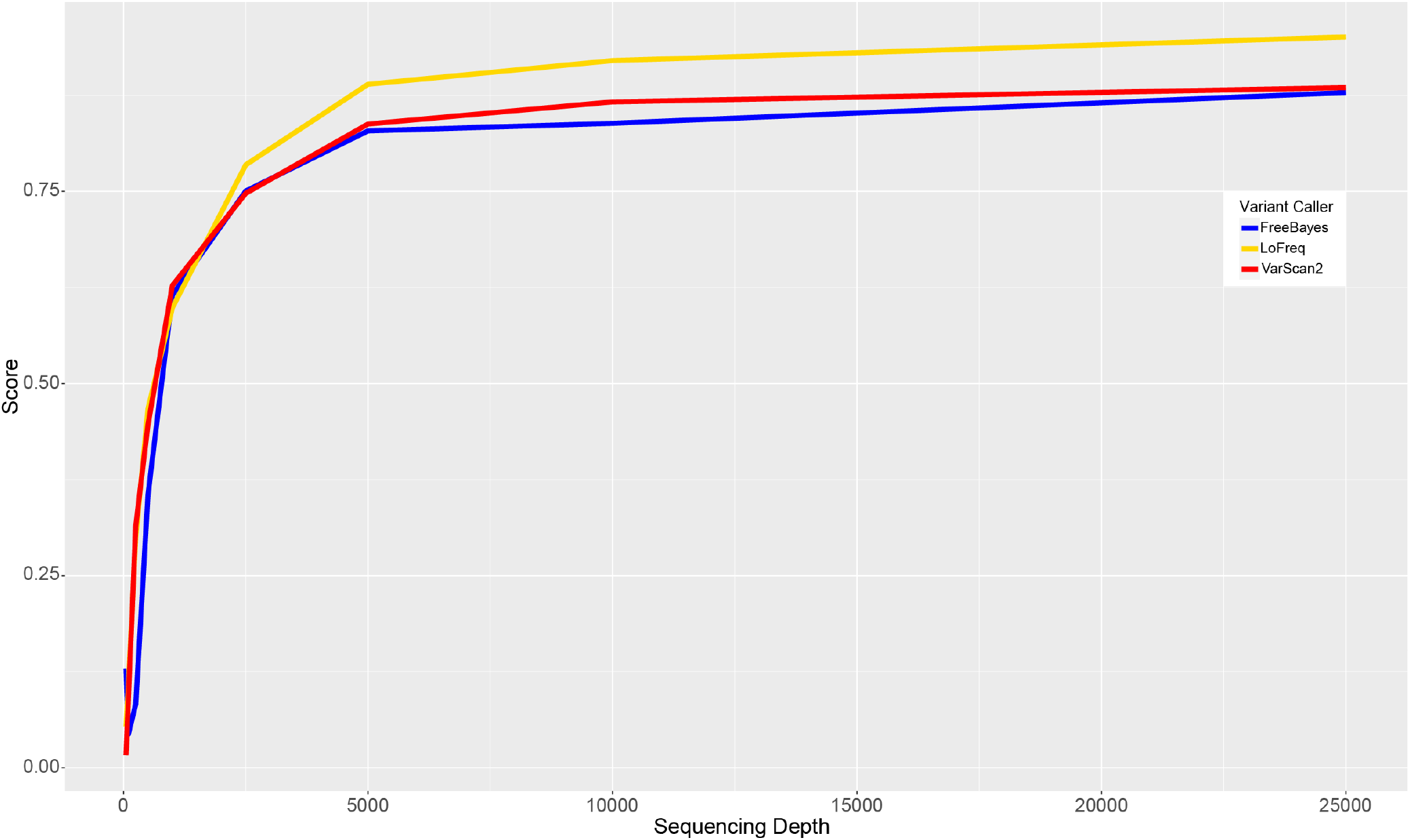
Results from simulated datasets using different variant callers. Nine read sets were generated *in silico* using GemSIM and the *M. myotis* mitochondrial genome, containing 500 known variants. The heteroplasmy detection pipeline was run on each dataset three times, using a different variant caller, with or without INDEL realignment and base quality recalibration as appropriate. The 27 resulting variant sets were compared to the known set for power, accuracy and false positive rate. A score was given to each set as (Power*Accuracy)*(1-False Positive Rate), and plotted against the expected coverage for the set. LoFreq was the best performing caller and was used for all variant calling on real data.

**Supplementary Figure 2:**
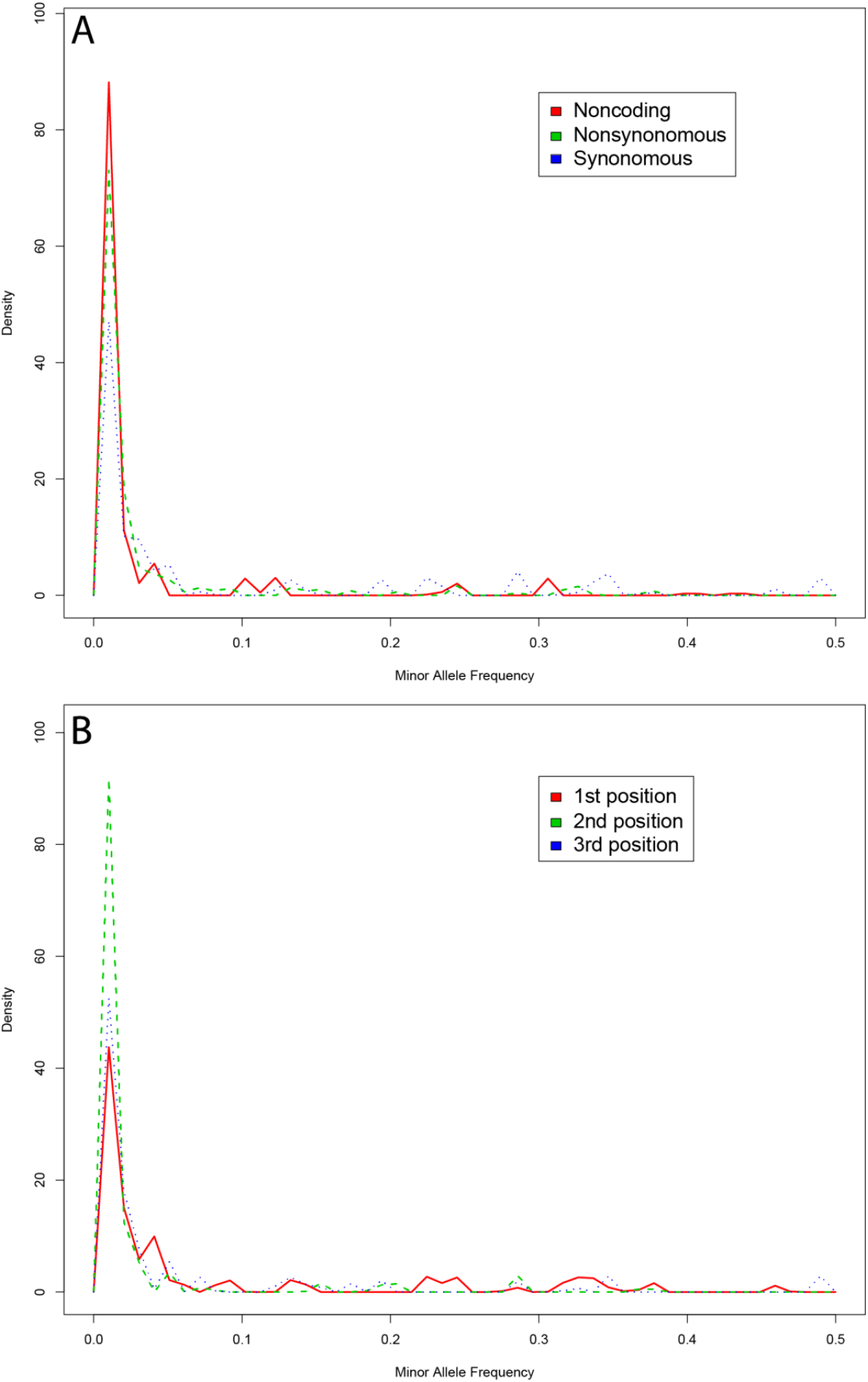
Frequency distributions and comparisons for different classes of heteroplasmies. **A)** Density plots of minor allele frequencies for noncoding, nonsynonymous coding and synonymous coding mutations. **B)** Density plots of minor allele frequencies for heteroplasmies at the first, second and third codon position.

**Supplementary Figure 3:**
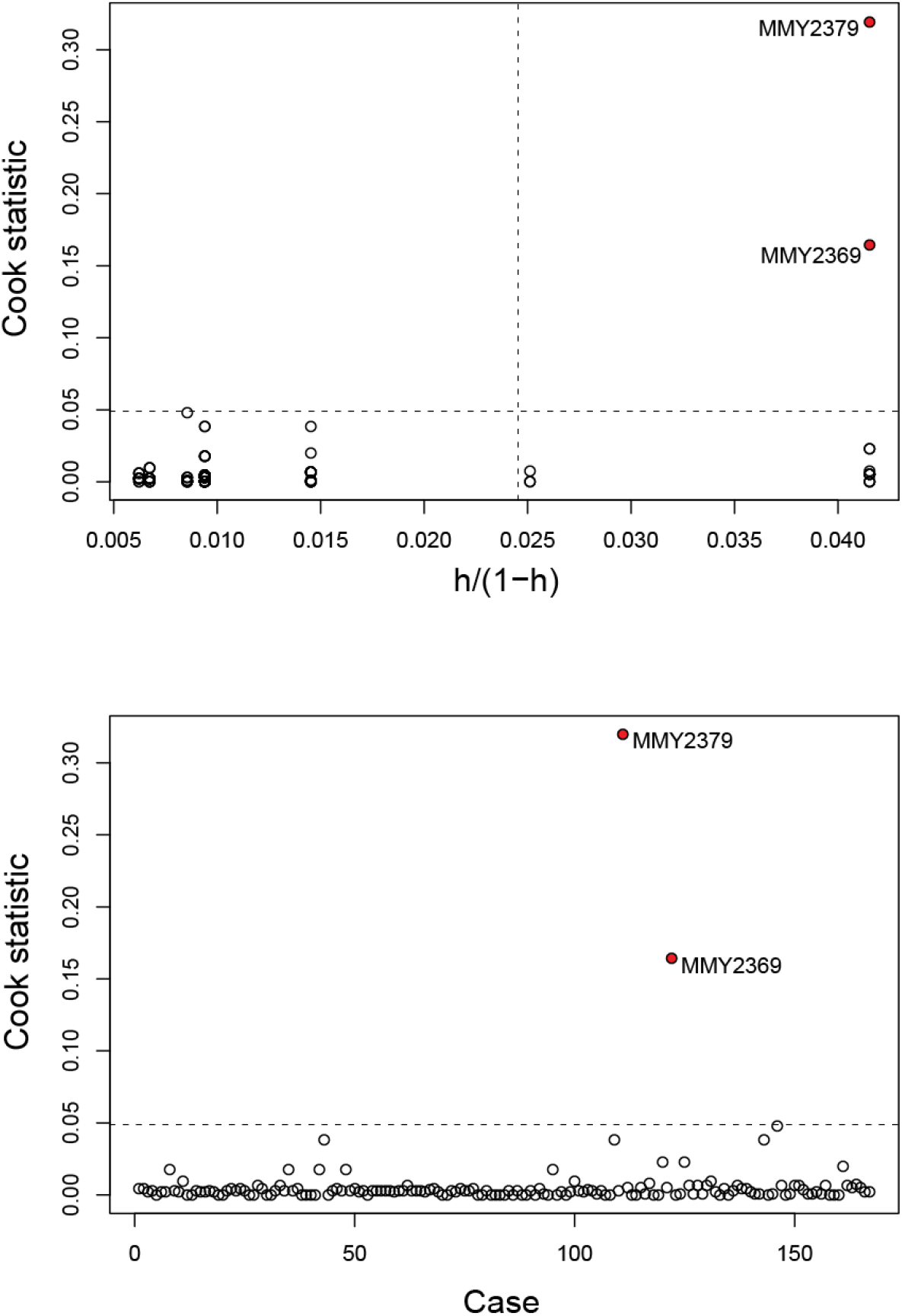
Cook’s Statistic predicting influential points in the Primary dataset. **A)** A plot of Cook’s statistics versus the standardised leverages for each point. The horizontal line is at 8/(n-2p) and the vertical line is at 2p/(n-2p) where n is the number of observations and p is the number of parameters estimated. Points above the horizontal line have a high influence on the model, and those to the right of the vertical line have a high leverage. The two outliers previously shown in Figure 4 are again indicated in red, with both above the horizontal and right of the vertical lines. **B)** A plot of the Cook’s statistics for each observation, order as they appear in the dataset. Again the two highly influential observations are coloured in red.

**Supplementary Figure 4:**
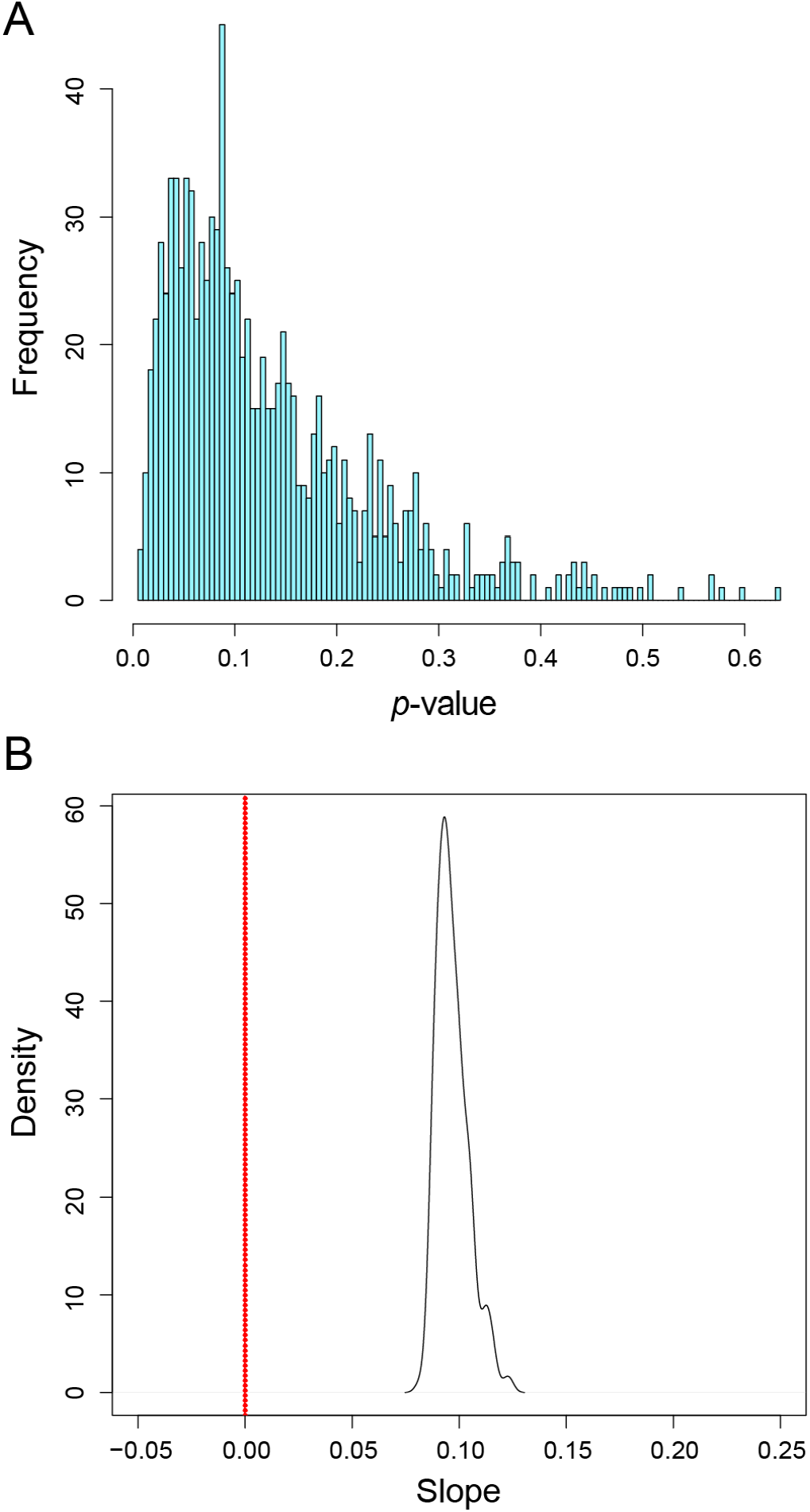
Test for the effect of sample selection on model fitting. Recaptured individuals of known age had more than one sample available for use in regression analyses. In the primary dataset the most recent, and thus oldest, sample from each recaptured was chosen. To ensure no bias was introduced by this selection, 1000 datasets were generated, randomly choosing which sample to use from the recaptured individuals, and then fitting the negative binomial model as before and estimating the significance and size of the age as the sole fixed effect. **A)** shows a histogram of p-values from 1000 models. 19.8% of models predicted a significant association between age and heteroplasmy **B)** shows a density plot of the slopes estimated from the 198 significant models, with a slope of 0 indicated by the red dotted line. The mean estimate was 0.0969 sites per year.

**Supplementary Figure 5:**
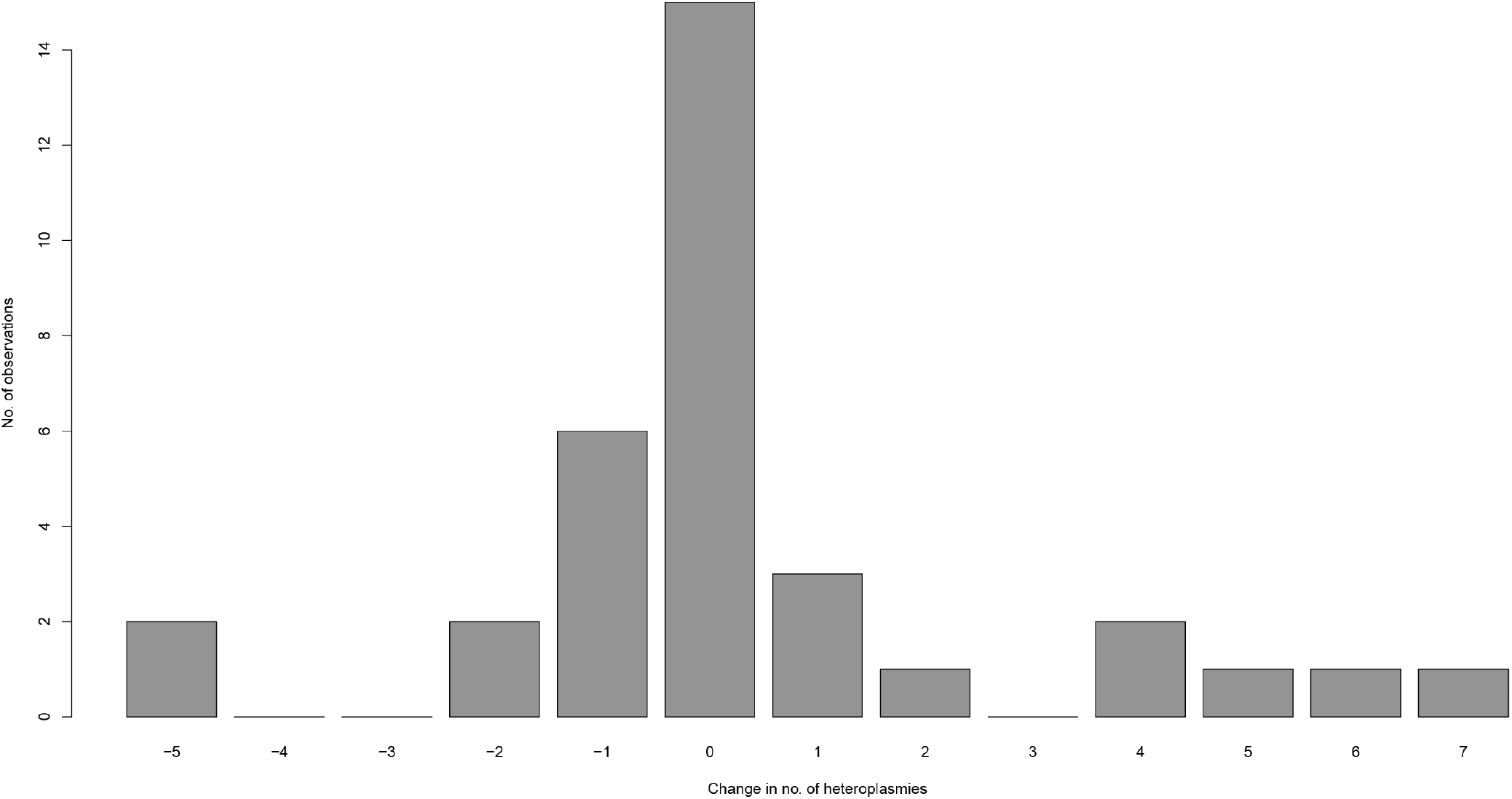
Change in the number of heteroplasmy between two consecutive years. Bar plot showing the number of times a change in heteroplasmy was observed, with respect to the size of the change, between two consecutive years. The mean change between any two years was 0.32, with median value of 0, though there were multiple observations of gains/loss of 5 or more heteroplasmies over a year.

**Supplementary Figure 6:**
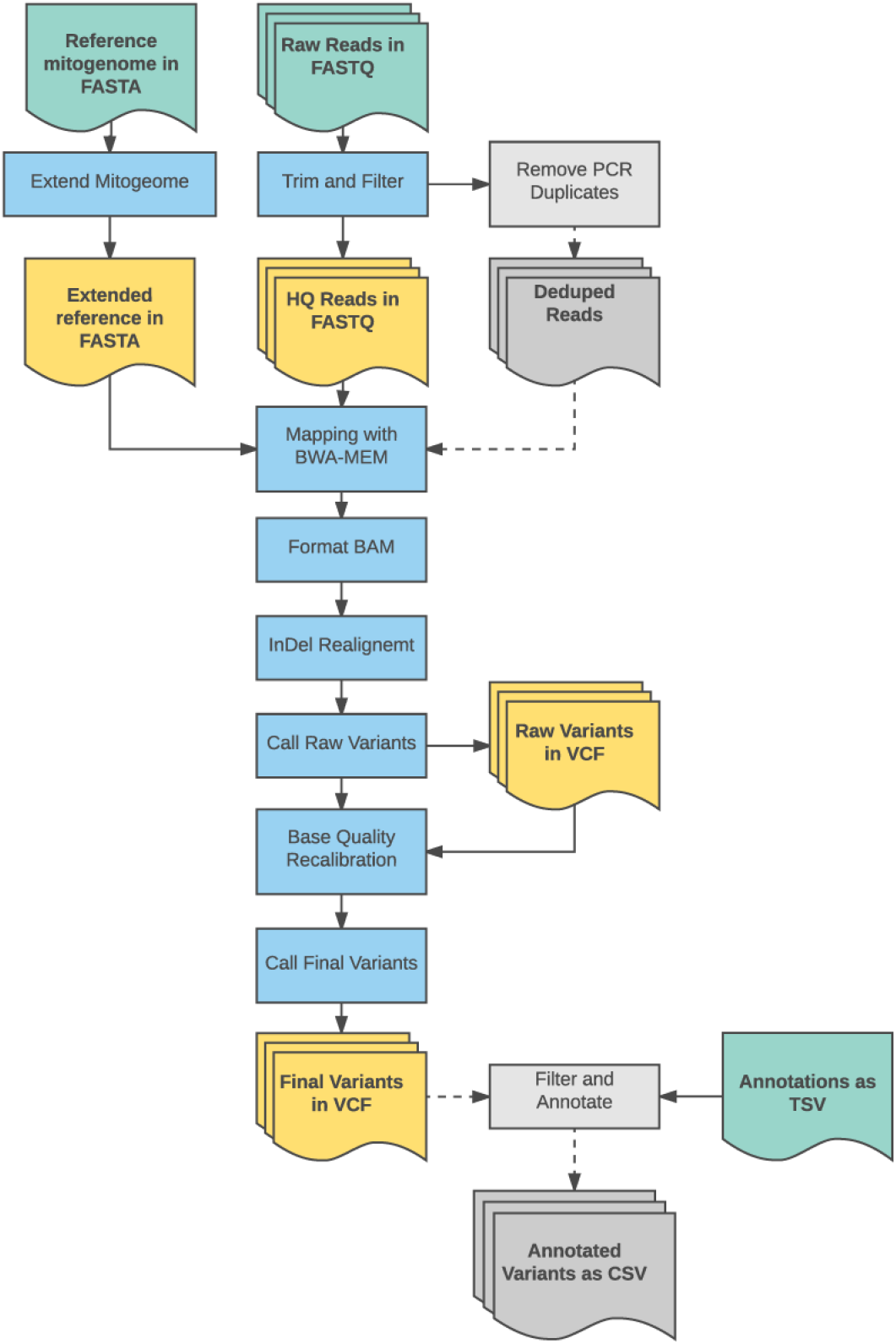
Overview of Heteroplasmy Detection Pipeline. Diagram depicting the bioinformatic workflow used in this project. Processes are depicted as rectangles, those in blue are mandatory, while those in grey are optional within the framework of the pipeline (duplicate removal and variant annotation). Documents are represented by rectangles with a wavy base. Those in green are input provided by the user (reference genome, annotations and sequence data), while the yellow documents are generated by default during analyses (extended reference, high quality reads, and variant calls) and grey documents can be generated but are optional (annotated variants). The pipeline can be parallelised, concurrently running a user defined number of jobs.

## Additional Files

Name: Additional File 1

Format: Microsoft Excel File (.xlsx)

Sheet 1: Simulation_results – Power, accuracy, false positive rate and score used to determine best variant caller from simulated datasets at different coverages

Sheet 2: Filtered_heteroplasmies – 254 annotated heteroplasmies discovered in this study

Sheet 3: Heteroplasmies_all_samples – Information for 232 samples including RFID, number of heteroplasmic sites and whether or not the sample was part of the primary dataset.

Sheet 4: Primary_dataset – Age, RFID and number of heteroplasmic sites for 167 samples used in the “Primary” dataset.

